# Pleiotropic and Non-redundant Effects of an Auxin Importer in Setaria and Maize^1^

**DOI:** 10.1101/2021.10.14.464408

**Authors:** Chuanmei Zhu, Mathew S. Box, Dhineshkumar Thiruppathi, Hao Hu, Yunqing Yu, Andrew N. Doust, Paula McSteen, Elizabeth A. Kellogg

**Affiliations:** Donald Danforth Plant Science Center, 975 North Warson Rd, Saint Louis, MO, 63132, USA; Department of Plant Biology, Ecology, and Evolution, Oklahoma State University, Stillwater, OK 74078, USA; Division of Biological Sciences, Interdisciplinary Plant Group, and Missouri Maize Center, University of Missouri, 301 Christopher Bond Life Sciences Center, Columbia, MO 65211, USA

## Abstract

Directional transport of auxin is critical for inflorescence and floral development in flowering plants, but the role of auxin influx carriers (AUX1 proteins) has been largely overlooked. Taking advantage of available AUX1 mutants in *Setaria viridis* and maize, we uncover previously unreported aspects of plant development that are affected by auxin influx, including higher order branches in the inflorescence, stigma branch number, and glume (floral bract) development, and plant fertility. However, disruption of auxin flux does not affect all parts of the plant, with little obvious effect on inflorescence meristem size, time to flowering, and anther morphology. In double mutant studies in maize, disruptions of *ZmAUX1* also affect vegetative development. A GFP-tagged construct of SvAUX1 under its native promoter showed that the AUX1 protein localizes to the plasma membrane of outer tissue layers in both roots and inflorescences, and accumulates specifically in inflorescence branch meristems, consistent with the mutant phenotype and expected auxin maxima. RNA-seq analysis finds that most gene expression modules are conserved between mutant and wildtype plants, with only a few hundred genes differentially expressed in *spp1* inflorescences. Using CRISPR-Cas9 technology, we disrupted *SPP1* and the other four *AUX1* homologs in *S. viridis*. SvAUX1/SPP1 has a larger effect on inflorescence development than the others, although all contribute to plant height, tiller formation, leaf, and root development. The AUX1 importers are thus not fully redundant in *S. viridis*. Our detailed phenotypic characterization plus a stable GFP-tagged line offer tools for future dissection of the function of auxin influx proteins.

**One sentence summary:** Mutations in a single auxin importer gene Spp1/SvAUX1 uncover broad and unexpected effects in nearly all aspects of the development of shoots, inflorescences, and flowers.

---

The plant hormone auxin is a mobile signal that is transported between cells by both influx and efflux proteins (Naramoto, 2017). It is involved in organ initiation and growth in all parts of the plant and is particularly well known for its effects on branching (Gallavotti, 2013; Taylor-Teeples et al., 2016; Naramoto, 2017; Olatunji et al., 2017; Korver et al., 2018). Efflux proteins, particularly homologs of PIN-FORMED1 (PIN1) (Petrásek et al., 2006; Balzan et al., 2014; Naramoto, 2017), have been studied extensively in many plant species, with particular attention in Arabidopsis, long the model of choice for studies of auxin function. As a result, much has been discovered about the flow of auxin out of cells (e.g. (Verna et al., 2019)) and how auxin gradients are established throughout the plant (e.g. (Heisler et al., 2005; Wang and Jiao, 2018) and many others).

In contrast, the flow of auxin into cells (auxin influx) has received much less attention, particularly in reproductive organs. In Arabidopsis single-gene mutants of any of the four auxin influx carriers (AUX1 and LAX1-3) have normal above-ground structures and higher order mutants affect only leaf phyllotaxis (Kleine-Vehn et al., 2006; Bainbridge et al., 2008; Peret et al., 2012; Swarup and Péret, 2012). Perhaps because of this subtle mutant phenotype, far less is known about influx than efflux, especially as regards vegetative and inflorescence development. Also the *AUX1/LAX* genes in Arabidopsis are more closely related to each other than any of them is to the *AUX1-like* genes known in grasses (Huang et al., 2017). This lack of one-to-one correspondence, in addition to the lack of a strong phenotype in Arabidopsis, prevents direct extrapolation from Arabidopsis to any monocot, particularly cereal crops and their relatives.

A recently identified mutation in an auxin influx carrier in the model grass *Setaria viridis*, SPARSE PANICLE1 (SPP1) (Huang et al., 2017), offers an opportunity to uncover novel aspects of auxin influx disruption. SPP1 is homologous to the maize protein ZmAUX1 and to the four Arabidopsis AUX1 proteins, but unlike in Arabidopsis, the *spp1* mutation (presumed to abolish gene function) causes an obvious defect in the inflorescence, thus providing a system in which the effects of disrupting influx are easily seen. SPP1 was named for the wide spacing of its primary inflorescence branches, and its role in auxin transport was supported by observation of clearly agravitropic roots (Huang et al., 2017). However, few other aspects of plant growth and development were considered in the original paper, including many that would be expected to require normal auxin transport. For example, the *S. viridis* inflorescence typically exhibits many orders of branches, some of which produce spikelets and others that end blindly (known as bristles; see (Doust and Kellogg, 2002)). Disruption of SPP1 should affect these higher order branches and the balance of spikelet-bearing branches and bristles, as well as other aspects of above-ground architecture such as tillering and relevant gene expression.

AUX1 mutants have been reported in other grasses (maize, rice, and Brachypodium) but these studies focused on roots (Yu et al., 2015; Zhao et al., 2015; Huang et al., 2017; van der Schuren et al., 2018), which were agravitropic in all species consistent with disruption of auxin pathways. In addition, the rice mutants had fewer lateral roots (Yu et al., 2015; Zhao et al., 2015), whereas the *S. viridis* mutants had a normal number (Yu et al., 2015; Zhao et al., 2015; Huang et al., 2017; van der Schuren et al., 2018). Neither Yu et al. (2015) nor Zhao et al. (2015) reported changes in the inflorescence in rice *OsAUX1* mutants. In *Brachypodium distachyon, bdaux1* mutants are sterile and some above-ground structures are affected, but the phenotypes are not described in detail (van der Schuren et al., 2018). Thus the role of AUX1 in above-ground development remains largely unexplored, especially in grasses and cereal crops.

Here we show that mutations in *SPP1* (=*SvAUX1*) and its homolog in maize affect many shoot phenotypes including development of the gynoecium and floral bracts (glumes); these are not side-effects of meristem size variation or differences in developmental timing. Based on the phenotypes of higher-order mutants involving all five *S. viridis* AUX1-like loci, we show that SPP1/SvAUX1 is not redundant with the other loci and is the major one controlling inflorescence architecture. *ZmAUX1*, investigated because of the wealth of auxin-related mutants in maize, enhanced the mutant phenotypes of several auxin pathway genes and revealed an unexpected enhanced effect on leaf number. In *S. viridis*, SPP1 was internally tagged, and localized to the plasma membrane of epidermal cells in inflorescence branch meristems and roots. Only a few hundred genes, including several known to be involved in inflorescence development, are differentially expressed between *spp1* and wildtype inflorescences, indicating highly specific changes in the transcriptome.

## RESULTS

### *spp1* affects tillering, inflorescence branching, gynoecium development, and root hair formation

Mutations in SPP1 affect many aspects of plant development having to do with growth and branching (Fig. 1; Table S1). In addition to the eponymous sparse panicle phenotype (Fig. 1A-1C), mutant plants were significantly shorter than wildtype (Fig. 1A, 1D) and produced more tillers (Fig. 1A, 1E). Mutant panicles were significantly longer than wildtype (Fig. 1B, 1C, 1F), but the increased length did not result in higher yield. Instead, mutants had fewer spikelets at maturity (Fig. 1G) and fewer of these were fully developed and fertile (Fig. 1H). The reduced number and fertility of spikelets was not caused by a developmental delay; the transition to reproductive growth and flowering in *spp1* mutant plants was only slightly later than in A10.1 (Fig. S1A, S1B), and barely significant. Fertile florets (upper lemma+palea) were significantly larger in the mutant (Fig. S1C) but percent germination did not differ (Fig. S1D). Culms (peduncles) were generally thinner in the mutant but overall culm anatomy was similar (Fig. S1E-S1G).

**Figure 1.**
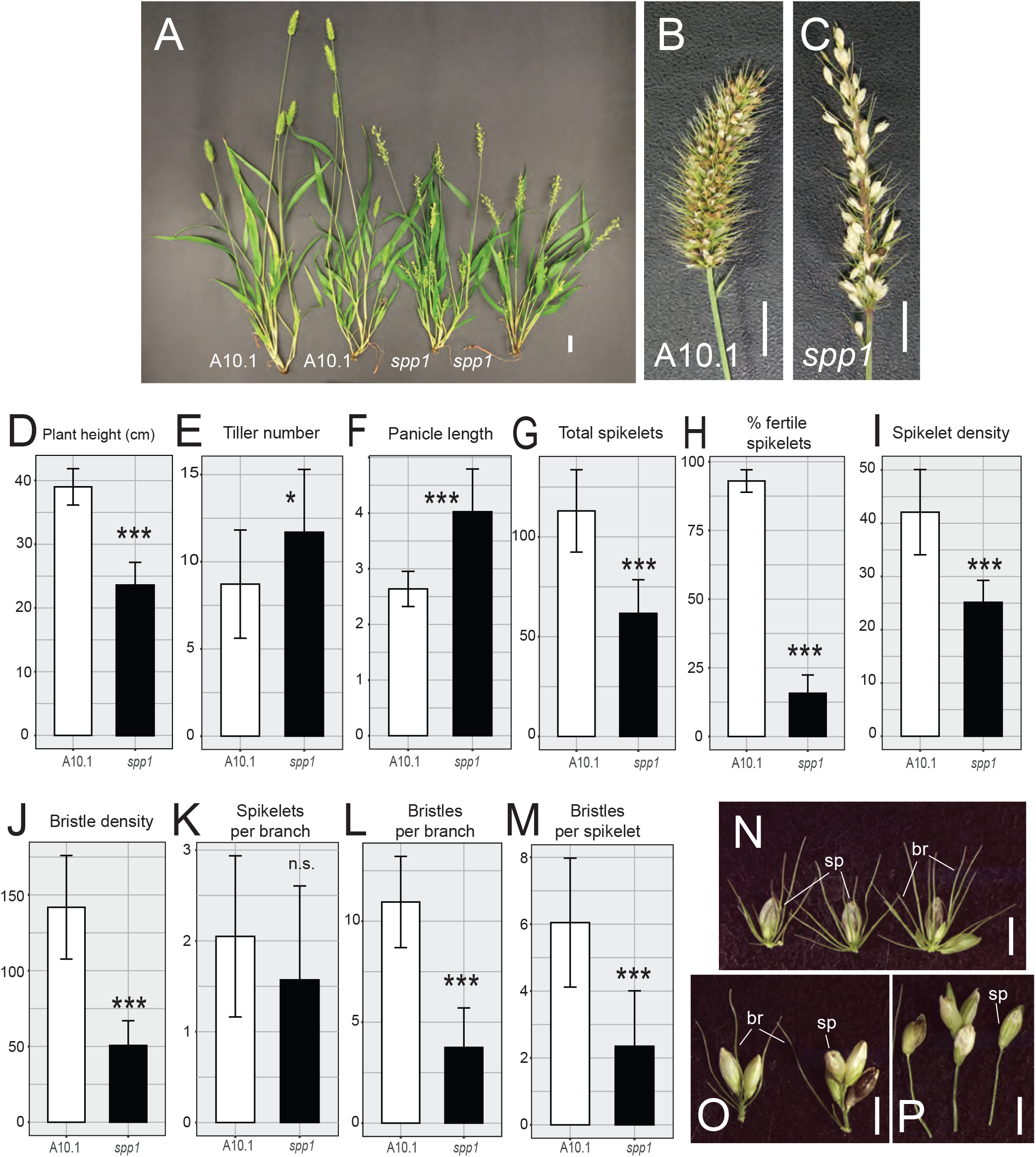
Phenotypes of *spp1* mutants. (A) Mature plants of wildtype (A10.1, left) and *spp1* mutants (right) at 22 DAS. Scale = 2 cm. (B, C) Mature panicles. (B) wildtype, (C) *spp1*. Scale = 1 cm. Brown or black spikelets contain fully developed seeds whereas whitish spikelets are often infertile. (D-M) Comparisons of trait values between wildtype (left bar, white) and *spp1* mutant (right bar, black) plants. Error bars, ± one standard deviation. Significance values determined by Welch’s t-test. = 0.01-0.05, *, <0.01, **, <0.001, ***, <0.0001. Values and sample sizes in Table S1. (D) Plant height (cm), (E) Tiller number), (F) Panicle length (cm), (G) Total number of spikelets, (H) Percent fertile spikelets, (I) Spikelet density (number of spikelets per cm), (J) Bristle density (number of bristles per cm), (K) Spikelets per primary branch, (L) Bristles per primary branch, (M) Bristles per spikelet (values from K divided by values from L), (N-O). Individual primary branches from wildtype (N) and *spp1* (O, P) mutants. Scale = 2 mm. sp, spikelet, br, bristle.

The lower density of spikelets and bristles (fewer of each per cm; Fig. 1I, 1J) could reflect a reduced density of primary branches (observed in early development; see next section) and/or a change in the numbers of spikelets and bristles per branch; the latter would indicate an effect of the mutation on secondary and higher order branches. In mutant panicles the primary branches have about the same number of spikelets as in wildtype (Fig. 1K, 1N, 1O), but significantly fewer bristles (Fig. 1L, 1N-1P) and therefore a lower ratio of bristles to spikelets (Fig. 1M-1P). In addition, about 15% of branches in *spp1* had one or a few spikelets at the terminus of a long branch without additional bristles, compared to <1% of A10.1 branches (Fig. 1P). Together these observations suggest that the *spp1* mutation affects both the formation of higher order branches and the specification of those branches as spikelets or bristles.

Floral morphology and early development are affected in *spp1* mutants and are likely to be at least partially responsible for the fertility defects of the mutant (Fig. 2; Table S1). At 18 days after sowing (DAS) when the anthers and gynoecium were first visible in both A10.1 and *spp1*, glumes in the wildtype were shorter than the flowers (Fig. 2A), whereas those in the mutants were unusually long, nearly enclosing the flowers (Fig. 2B). In addition, the mutants had fewer branches, bristles, and spikelets at this stage, consistent with the reduced number of bristles per spikelet at maturity (Fig. 1M). Gynoecium formation was also abnormal in *spp1*, with mutants often having fewer than two styles, the normal number in wildtype (Fig. 2C-2G). Stigmas in *spp1* plants, when present, were significantly less branched than in wildtype (Fig. 2H-2J); Table S1). Other than the abnormal glume and gynoecium development, all spikelets in both genotypes had the expected number of glumes (two) and florets (two), with lemmas, paleas, lodicules and stamens developing apparently normally in both mutant and wildtype plants (Fig. 2D-2F).

**Figure 2.**
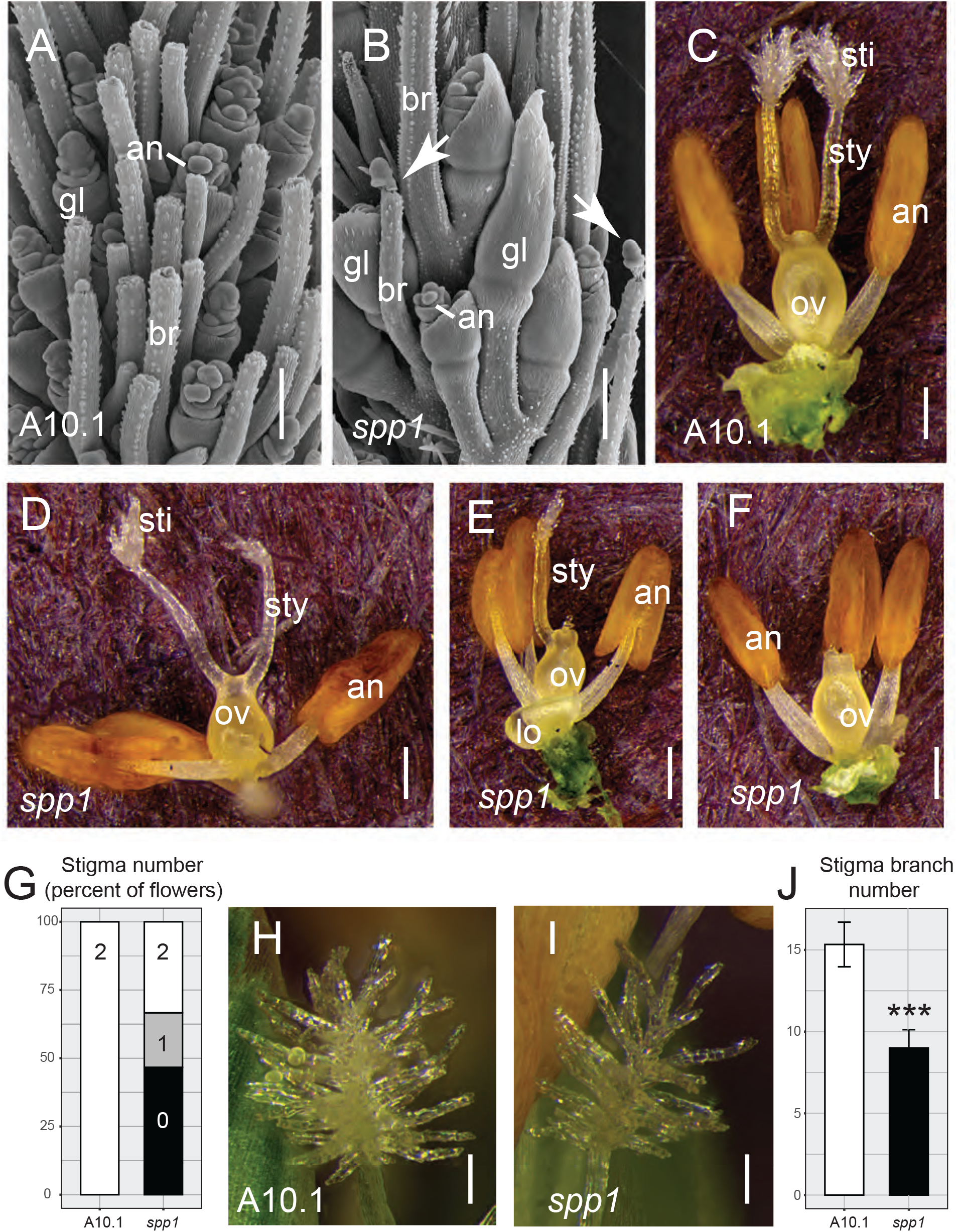
Floral phenotypes of *spp1*. (A, B) SEM images of developing spikelets and bristles at 18 DAS. (A) wildtype (A10.1), (B) *spp1*. Arrows show un-detached meristems on bristle tips. Scale = 200 μm. (C-F) Reproductive organs in wildtype (C) and *spp1* (D-F) mutant florets, showing abnormal development of stigmas and styles in the mutants. Scale = 250 μm. (G) Bar graph showing percentage of florets with 0, 1 or 2 stigmas in wildtype (left) and *spp1* (right). (H, I) stigmas from wildtype (H) and *spp1* florets (I). Scale = 100 μm. (J) stigma branch number counted from one side of the stigma on the focal plane in wildtype (left, white) and *spp1* (right, black) plants. Error bars, ± one standard deviation. significance values as in Figure 1. Values and sample sizes in Table S1. an, anther; br, bristle; gl, glume; lo, lodicules; ov, ovary; sti, stigma; sty, style. Image (A) reproduced with permission from Zhu et al. (2018).

Neither primary root length nor lateral root number was obviously altered in *spp1* (Fig. S1N), but root hair density was significantly lower on both primary and lateral roots in *spp1* compared to A10.1 (Fig. S1H, S1I, S1L, S1M; Table S1). In addition, the distance from root tip to the first root hair initiation site was significantly longer in mutant roots (Fig. S1J, S1K).

By applying synthetic auxins to roots, we showed that SPP1 could potentially function in auxin import. In response to a mock auxin treatment, *spp1* roots were agravitropic (Fig. S2A, S2B), as expected (Huang et al., 2017), and had fewer root hairs than wildtype (Fig. S2G, S2H). 2,4-Dichlorophenoxyacetic acid (2, 4-D), which requires auxin importer proteins to move into the cells, could not rescue the mutant phenotypes in roots, consistent with our hypothesis that SPP1 is a *bona fide* auxin importer (Fig. S2C, S2D, S2I, S2J, S2M). In contrast, the lipophilic auxin 1-Naphthaleneacetic acid (NAA), which can diffuse freely across the plasma membrane, restored both the gravitropic response of *spp1* roots (Fig. S2E, S2F) and also the normal density of root hairs (Fig. S2K-S2M).

### SPP1 controls inflorescence branch initiation, elongation, and identity, but not meristem size

To explore whether the sparse panicle phenotype in *spp1* resulted from branch initiation defects linked to abnormal meristem size, we imaged early inflorescence development with scanning electron microscopy (SEM)(Fig. 3A-3N; Table S2). Meristem height (the vertical distance from the uppermost branch primordium to the apex of the meristem) dropped significantly between 11 and 12 DAS and again between 12 and 13 DAS, but wildtype and mutant inflorescences did not differ at any stage of development (Fig. 3O). Meristem width was unchanged in either genotype over 10-12 DAS, then dropped significantly in both genotypes between 12 and 13 DAS (Fig. 3P); by 14 DAS, *spp1* inflorescences were wider than those in wildtype (Fig. 3P). Overall length of inflorescences before 14 DAS scarcely differed between *spp1* and wildtype (Fig. 3A-4N, 3Q), indicating that the length difference at maturity was established later in development and probably reflected rachis elongation rather than branch initiation. By 12 DAS, primary branch number in *spp1* was significantly lower than in A10.1, whether counting branches per vertical row (Fig. 3R), or all visible branches on one side of the inflorescence (Fig. 3S). In contrast to A10.1, which produced primary branch meristems in a spiral pattern around the inflorescence meristem (Fig. 3A-3E, 3K, 3L), *spp1* often failed to initiate a branch meristem or produced unusually large primary branch meristems (Fig. 3F-3J, 3M, 3N). While primary branch meristems produced distichous secondary branch meristems in A10.1 (Fig. 3C-3E, 3K, 3L), secondary branches often initiated asymmetrically in *spp1* (Fig. 3H-3J, 3M, 3N).

**Figure 3.**
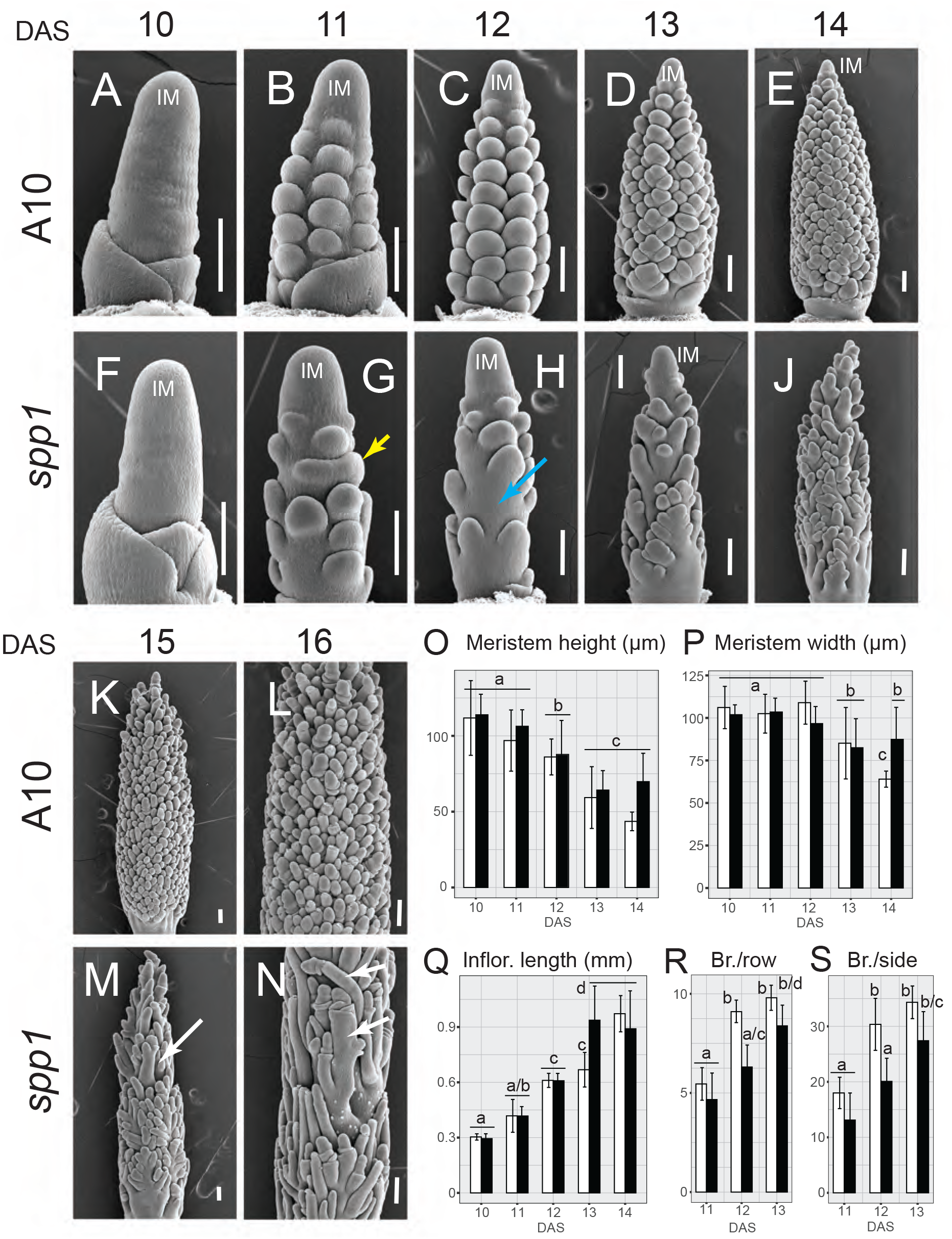
Early inflorescence development of *spp1*. (A)-(N) SEM images of wildtype (A10.1) (A-E, K, L) and *spp1* (F-J, M, N) inflorescences at 10-16 DAS (left to right, one picture for each stage, respectively). Yellow arrow, fused primary branch meristems; blue arrow, failed initiation of primary branch meristem; white arrows, elongated branch primordia. (O-S) Comparisons of wildtype (white) and *spp1* (black) inflorescences as measured from SEM photos. (O) meristem height and (P) meristem width (μ) at 10-14 DAS. (Q) Inflorescence length (mm) at 10-14 DAS. (R, S) Number of primary branch meristems per vertical row (R) and the total number visible from one side of the inflorescence (S). Error bars, ± one standard deviation. Significance values determined by ANOVA; p values indicated as in Figure 1. Bars with the same letter are not significantly different. Values and sample sizes in Table S2. IM, inflorescence meristem.

*spp1* was defective in branch elongation and meristem fate determination. Branch primordia in *spp1* elongated more than those in A10.1 (Fig. 3A-3N). While most bristles in A10.1 had lost their meristematic tip completely by 16 DAS (Fig. 3L), bristles often retained their meristem in *spp1* (Fig. 3N) even at 18 DAS (Fig. 2A, 2B).

### The *Spp1* ortholog in maize, *ZmAux1*, enhances effects of auxin-related genes

Because *S. viridis* lacks a set of auxin-related mutants, we used maize to test genetic interactions of AUX1 with other loci. The mutant for the *SPP1* ortholog in maize, *zmaux1*, produced fewer branches in the tassel and fewer spikelets per row in the ear and tassel compared to the wildtype (W22 inbreds) and heterozygous controls (Table S3), a phenotype analogous to that in *S. viridis* (Fig. 4A-4G). Also like *S. viridis*, the mutation had no obvious effect on inflorescence meristem sizes (Fig. 4A-4F). Spikelets in *Zea* occur in pairs, with a pair generally interpreted as a short lateral branch (Vollbrecht et al., 2005; Whipple, 2017). Therefore if *zmaux1* affects higher order branches in the inflorescence, it should affect whether both members of the pair initiate and indeed *zmaux1* showed more single and fewer paired spikelets in both ear and tassel (Fig. 4A-4F, 4H; Table S3). The tips of *zmaux1* ears were often elongated as were some individual spikelets themselves (Fig. 4E), similar to the spikelet-tipped bristles in the *spp1* mutant. Thus *SPP1* controls branch initiation, elongation and fate determination, but not inflorescence meristem size, in both *S*.*viridis* and maize.

**Figure 4.**
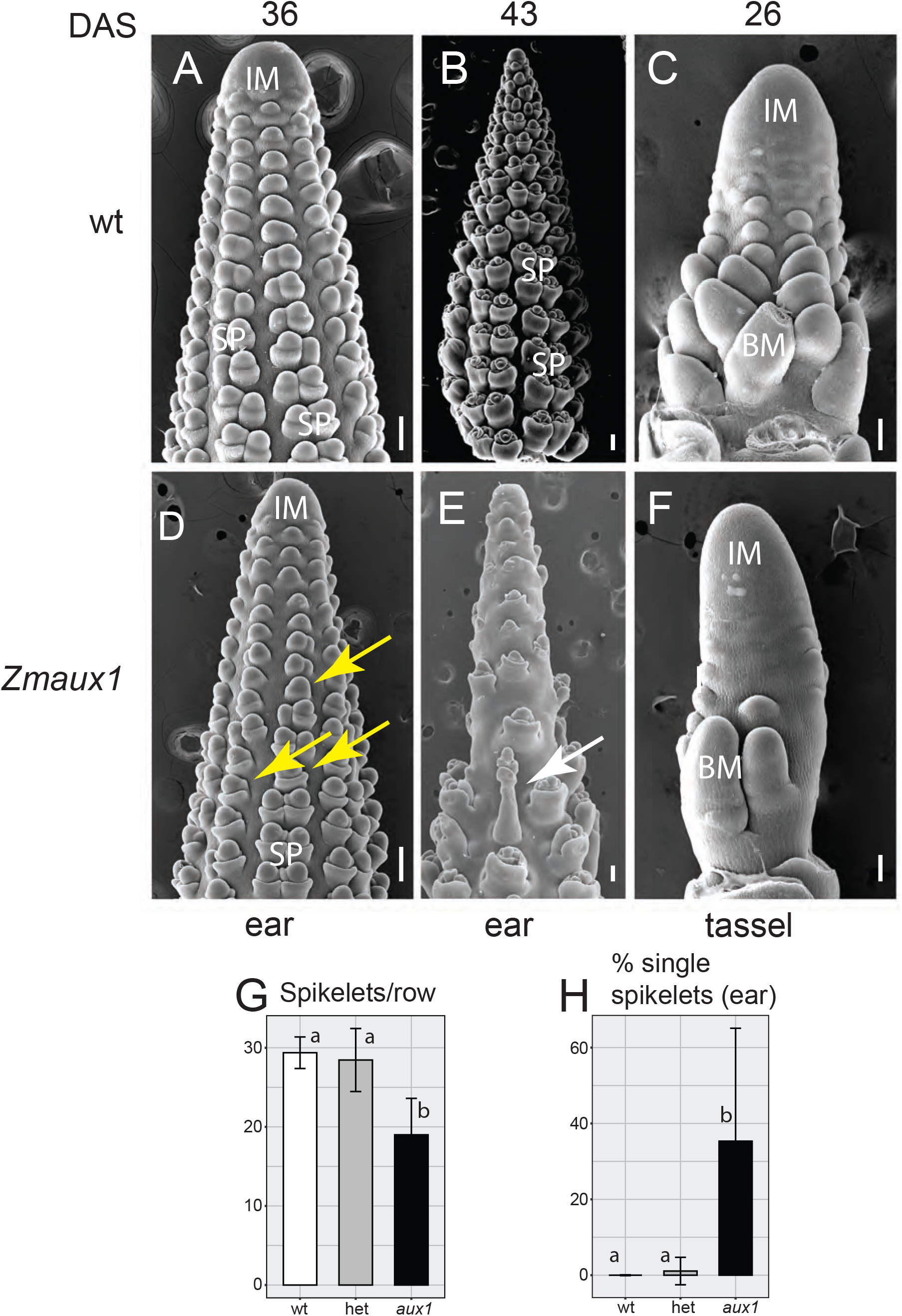
Early ear and tassel inflorescences of *zmaux1*. (A)-(F) SEM images of wildtype (W22) (A-C) and *zmaux1* (D-F) inflorescences. (A, B) are heterozygous wt; *Zmaux1* (C) is homozygous wildtype. Ears (A, B, D, E) at 36 (A, D) and 43 DAS (B, E). Tassel at 26 DAS (C, F). Yellow arrows, single spikelets. White arrow, elongated spikelet. (G) Number of spikelets per vertical row in the ear in wildtype (white), heterozygote (gray) and *zmaux1* (black) plants. (H) Percentage single spikelets in ear. Colors as in (G). Error bars, ± one standard deviation. Significance values determined by ANOVA; p values as in Figure 1. Bars with the same letter are not significantly different. Values and sample sizes in Table S3. Scale = 100 μm. BM, branch meristem; IM, inflorescence meristem; SP, spikelet pair.

We crossed three well-characterized auxin mutants in maize to *zmaux1*, guided by the presumed pathway shown in Fig. 5A based on their biochemical functions. These included an auxin biosynthesis mutant (*vanishing tassel 2* (*vt2*), encoding a grass-specific tryptophan aminotransferase) (Phillips et al., 2011), a regulator of auxin efflux (*barren inflorescence 2* (*bif2*), encoding a serine/threonine kinase co-orthologous to PINOID in Arabidopsis) (McSteen et al., 2007; Pressoir et al., 2009), and an auxin signaling protein (*Barren inflorescence 4* (*Bif4*), encoding an AUXIN/INDOLE-3-ACETIC ACID (Aux/IAA) protein) (Galli et al., 2015).

**Figure 5.**
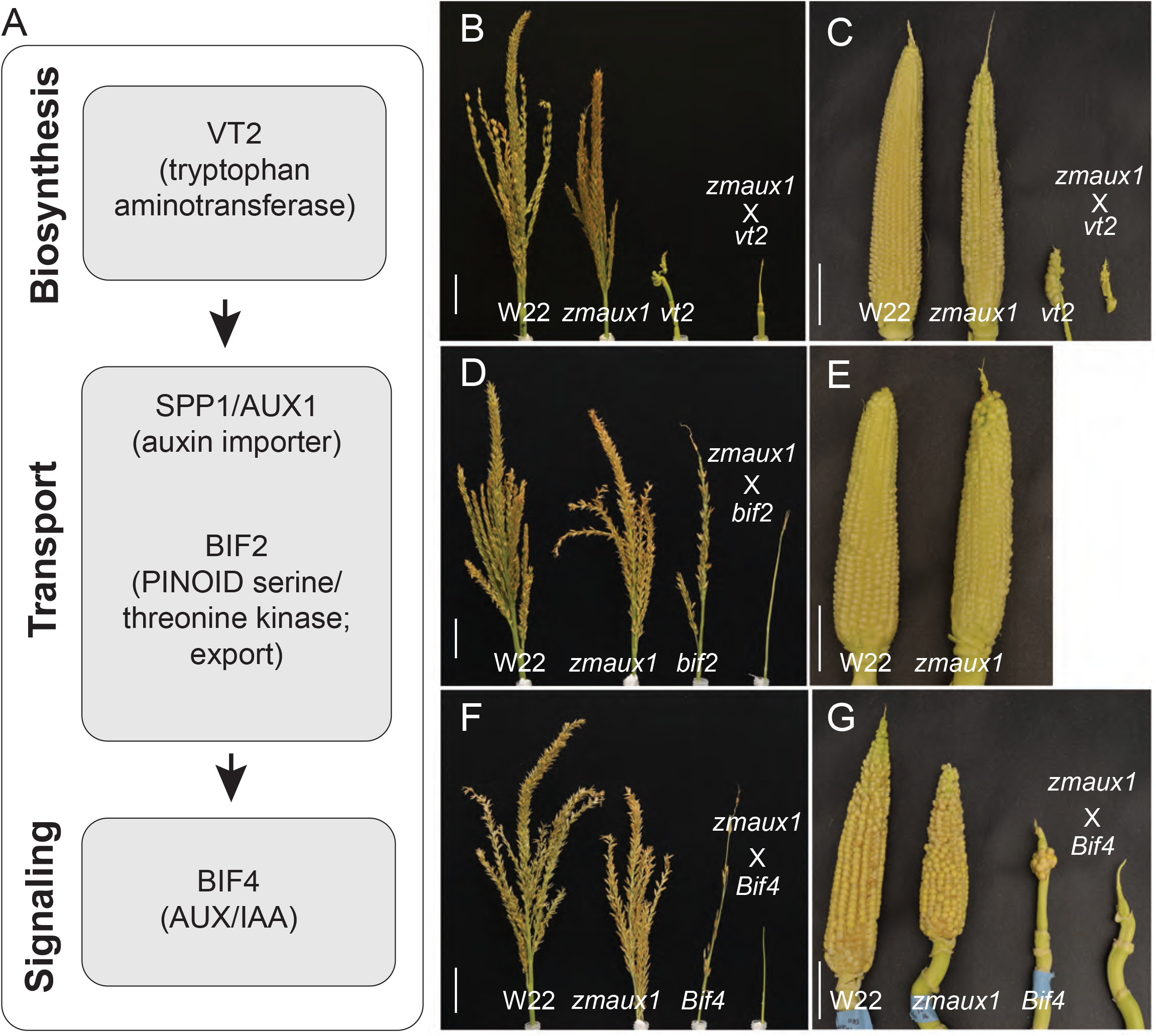
Auxin double mutant analysis in maize. (A) Model showing hypothesized relationship of classic genes involved in auxin biosynthesis, transport and signaling, based on information from the literature regarding function. (B, D, F) tassels and (C, E, G) ears from F2 progeny of crosses between *zmaux1* and *vt2* (B, C), *bif2* (D, E), and *Bif4* (F, G). Genotypes in each panel are, left to right, wildtype (WT), *zmaux1*, classical mutant, and double mutant. Most *bif2* and *zmaux1bif2* mutants fail to produce ears. Scale = 5 cm.

Plants with the mutant allele *zmaux1* had reduced branching in the ear and tassel in all three mutant families (*vt2, bif2, Bif4*)(Figs. 5B-5G, S3-S5). Kernel number, reflecting the total number of spikelets and hence the total number of branches, was also significantly reduced by the *zmaux1* single mutant in all mutant families, although traits that might contribute to total kernels (ear row number, spikelets per row) were not significantly lower in all cases, probably due to small sample size (Table S3; Figs. S3D, S4D, S5D). Number of tassel branches was also significantly lower in all cases, but the number and density of spikelets on the main spike of the tassel was not always affected. In contrast, tassel length, height of the flag leaf, and total number of leaves were not significantly different for *zmaux1* mutants (Table S3; Figs. S3-S5).

The effect of the double mutants on inflorescence characteristics is consistent with what we know about the function of the underlying genes. The locus defective in auxin biosynthesis, *vt2*, almost completely abolished branching in both the tassel and ear and suppressed growth of the tassel, thereby completely obviating any effect of *zmaux1. vt2* single mutants were indistinguishable from *zmaux1;vt2* double mutants for these traits (Fig. 5B, S3B-S3G). Likewise, BIF2 phosphorylates the auxin efflux carrier ZmPIN1 and its mutation blocks inflorescence branching, presumably by preventing auxin efflux (Skirpan et al., 2009). *bif2* single mutants were also indistinguishable from the *zmaux1;bif2* double mutant for the same branching traits as *vt2* (Fig. S4B-S4G). *Bif4* encodes a protein involved in auxin signaling and creates a less severe defect in branching than *vt2* and *bif2* (Fig. S5). The *Bif4* mutant phenotype is significantly enhanced in the *zmaux1;Bif4* double mutant for kernel number, tassel branch number, and density of spikelets on the main spike of the tassel, although the effect on ear row number, spikelets per row, and spikelets on the main spike was non-significant (Fig. S5B-S5G).

The double mutants had an unexpected effect on vegetative characteristics. The *vt2* and *bif2* mutations led to slight but non-significant reductions in flag leaf height and leaf number, an effect that was clearly enhanced by *aux1*; the phenotype of the double mutants *zmaux1;vt2* and *zmaux1;bif2* was significantly more severe than either single mutant (Fig. S3I, S3J, S4I, S4J). The vegetative traits in the *Bif4* family were even more striking in that neither *zmaux1* nor *Bif4* single mutants had a significant effect but leaf number and height of the flag leaf were reduced in double mutants (Fig. S5I, S5J). The synergistic effect in double mutants involving all three auxin-related genes indicated that *zmaux1* does indeed function in the auxin pathway, and moreover that auxin import has a role in normal leaf organogenesis.

### SPP1 localizes to epidermal cells in branch meristems in the inflorescence

SPP1 was localized in the *S. viridis* inflorescence using a translational fusion with a green fluorescent protein (GFP) fused to SPP1 (SPP1~iGFP) in an internal facing (cytoplasmic) N-terminal hydrophilic loop of SPP1 (Fig. S6A). We initially placed *SPP1~iGFP* under a constitutive promoter (*proPvUBI1::SPP1~iGFP*) to check its integrity with transient expression assays in tobacco leaves. SPP1~iGFP localized preferentially to a thin line at the periphery of epidermal cells, consistent with plasma membrane (PM) localization (Fig. S6B-S6D).

SPP1~iGFP localization is consistent with its routing through the secretory pathway to the plasma membrane as well as the nuclear membrane. Using tissue culture transformation, we introduced our *SPP1~iGFP* construct driven by its native promoter (*proSPP1::SPP1~iGFP*) to *spp1* mutants, validated three independent events by PCR genotyping, and selected one containing an expressed transgene (*spp1_T*) for further characterization (Fig. S7A, S7B). Confocal imaging in the T3 generation showed that in developing inflorescence, emerging leaves, and roots, GFP signals were mostly on the cell periphery of outer epidermal layers (Fig. 6A-6F, S6E-S6J). SPP1~iGFP in leaves colocalized with FM4-64, a marker of the plasma membrane, confirming that the peripheral location of the signal indeed came from the membrane (Fig. 6D-6F). SPP1~iGFP was also visible in a fine perinuclear line, likely to be the nuclear membrane (Fig. 6C, S6H, S6I), and in transcellular strands extending from the nucleus to the plasma membrane (Fig. 6C).

**Figure 6.**
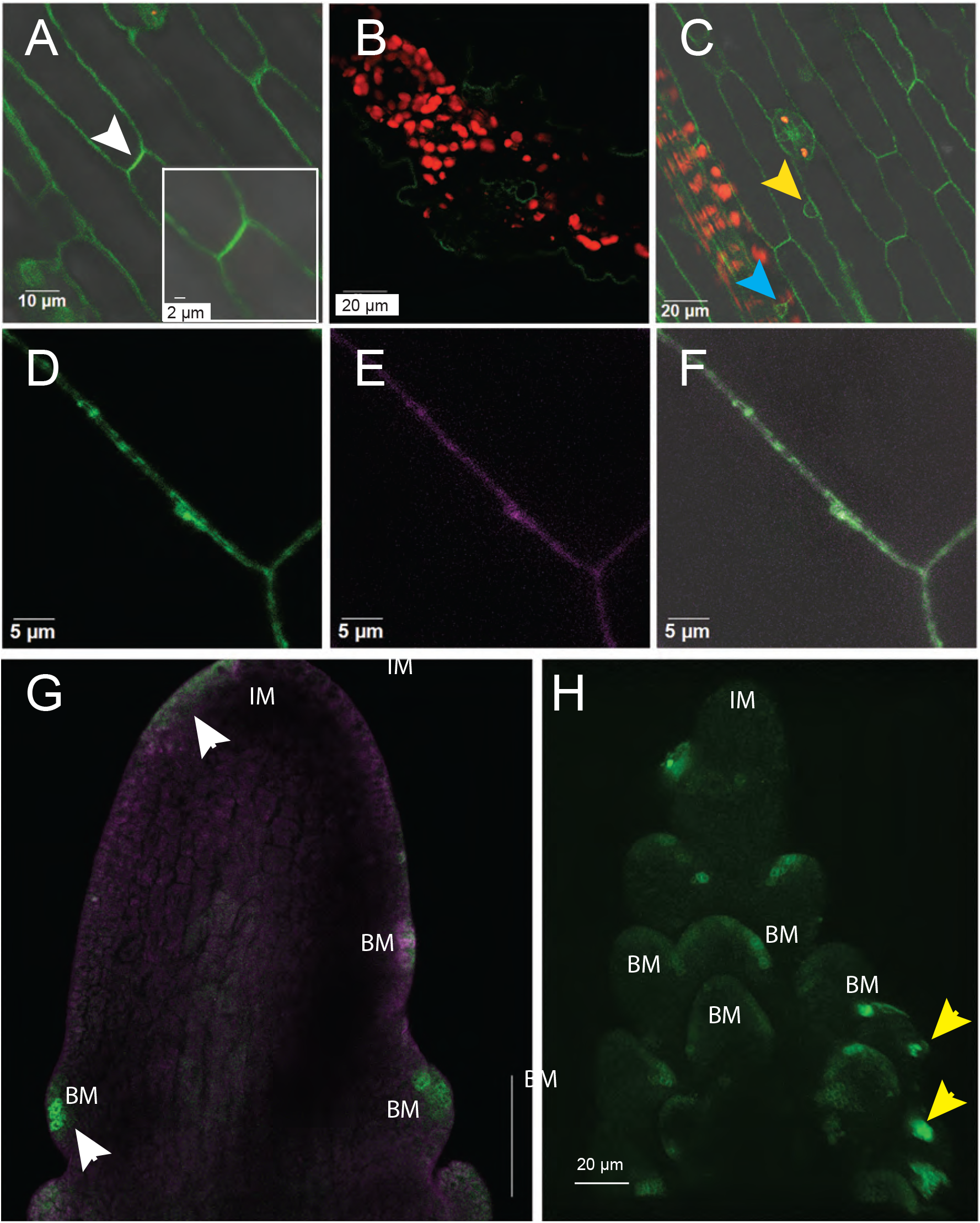
Expression pattern and subcellular localization of SPP1~iGFP in *S. viridis*. (A-F) Localization of SPP1~iGFP in stably transformed *S. viridis* leaves at 8 DAS. (A) Leaf surface showing fluorescent signals on the plasma membrane (PM). Strongest signals on PM may indicate weak polar localization (white arrowhead). (B) Leaf cross section showing SPP1 expression in epidermis and veins. (C) Leaf showing weak GFP signals on the transcellular strands (cyan arrowhead) extending from nucleus to PM, and around the nuclear membrane (yellow arrowhead). Red, chlorophyll autofluorescence. (D-F) Leaf cells expressing SPP1~iGFP (D; green), counterstained with FM4-64 (E; magenta), visible as a thin line on PM. Overlay (F) merges (D) and (E). (A, C-F) are single confocal sections; (B) is a projection of several sections. Scales as noted on images. (G, H). Localization of SPP1~iGFP in stably transformed *Setaria* inflorescences at 11 DAS. (G) Expression of SPP1~iGFP fusion protein appears in initiation sites of primary branch meristems along inflorescence flanks (white arrowheads). IM lacks fluorescent signals. See also Supplemental Video 1. (H) A single epidermal confocal focal plane from Video S2 showing epidermal enrichment of SPP1~iGFP expression in meristems of elongating primary branches. A few secondary branches also express SPP1~iGFP (yellow arrowheads). Merged image of green (GFP signals) and magenta (for FM4-64 signals) channels. IM, inflorescence meristem; BM, branch meristem.

SPP1~iGFP appeared at the presumed initiation site for all branch meristems, both primary and secondary. SPP1~iGFP was visible on the side of IM at about 1-2 days after IM formation (Fig. 6G, Video S1), likely marking the position of the youngest primary branch meristem. Expression decreased or disappeared in older IMs and older branch meristems (Fig. 6H, Video S2). Consistent with the expression pattern, SPP1~iGFP partially rescued defects in *spp1* inflorescences (Fig. S7C-S7I; Table S4), although the effect of the transgene was only significant for panicle length (Fig. S7F) and spikelets per primary branch (Fig. S7H). It also rescued the agravitropic root phenotype of *spp1* (Fig. S7J, S7K). Partial rescues are common and are thought to reflect the failure of the transgene to completely mimic endogenous gene expression (Stam et al., 1997).

#### *SPP1* affects expression of inflorescence developmental genes

We used RNA-seq to compare gene expression in A10.1 and *spp1* inflorescences at 10, 12 and 14 DAS (see Fig. 3, Tables S5-S7); transcripts were clustered with WGCNA (Langfelder and Horvath, 2008). Among the 10,434 transcripts in the analysis, we identified seven co-expression modules in A10.1 inflorescences and ten in *spp1* (Fig. S8). None of the modules was genotype-specific and most were strongly preserved between genotypes (Fig. S9). For example, the largest module in A10.1 (turquoise) included 6650 transcripts with low expression at 10 DAS, moderate at 12, and high expression at 14 DAS; 5,571 of these transcripts fell into either the turquoise or blue modules in *spp1*, which showed a similar overall pattern (Fig. S8, S9B). Most GO terms were comparable between the two genotypes, but the terms “cellular response to auxin stimulus”, “response to gibberellin” and “regulation of abscisic acid-activated signaling pathway” showed differential enrichment (Fig. S10).

Consistent with the high conservation of the WGCNA expression modules, relatively few transcripts were differentially expressed between A10.1 and *spp1*. At 10 DAS, before the mutant phenotype was visible, only 166 genes were differentially expressed, 57 of which differed more than two-fold (Fig. 7A, Table S7). At 12 and 14 DAS, still only a few hundred genes were differentially expressed (Fig. 7A, Table S7), with slightly more downregulated than upregulated in the mutant compared to wildtype.

**Figure 7.**
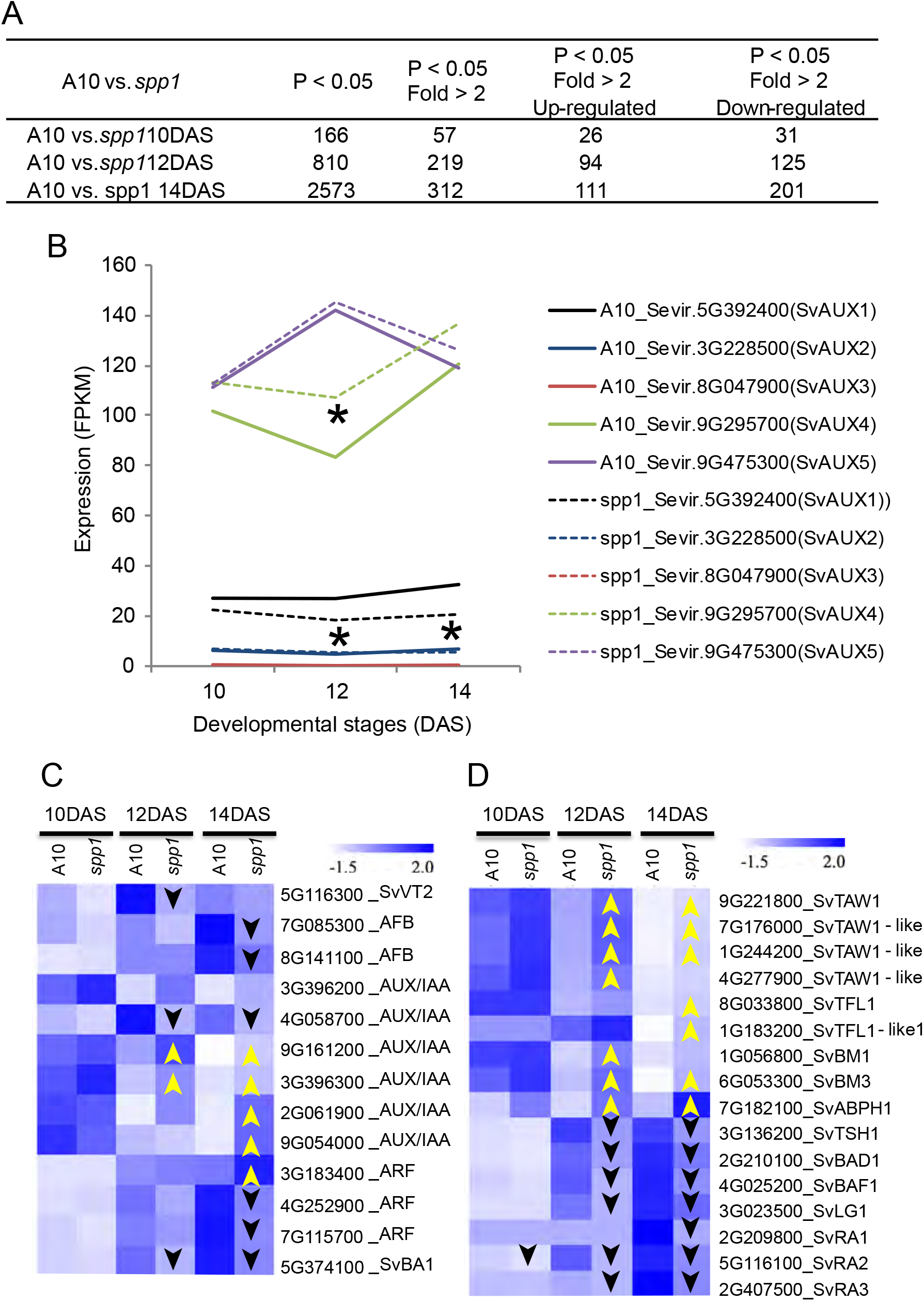
Differentially expressed genes in *spp1* inflorescences at 10, 12 and 14 DAS. (A) Numbers of genes that are differentially expressed, upregulated or downregulated between wildtype (A10.1) and *spp1* at each time point. (B) Expression of the five auxin influx carrier genes in *S. viridis* in wildtype and *spp1* inflorescences. (C) Heat map comparing expression of selected auxin-pathway related genes in wildtype (A10.1) and *spp1* inflorescences. (D) Heat map of selected differentially expressed genes involved in inflorescence branching. Yellow upward pointing arrows and black downward pointing arrows indicate upregulation and downregulation, respectively, compared to A10.1 at the same developmental stage.

We investigated the expression of *SPP1* and its four homologs, *SvAUX2-SvAUX5* (Fig. 7B); *SvAUX1* is *SPP1. SvAUX1* expression in *spp1* was significantly reduced at 12 and 14 DAS compared to that in A10.1 (Fig. 7B), as shown previously with qRT-PCR (Huang et al., 2017). At all three time points of both genotypes, expression of *SvAUX2* was several-fold lower than that of *SvAUX1* and *SvAUX3* was scarcely expressed at all (FPKM values <1 for all samples; Table S6). *SvAUX4* and *SvAUX5* were more highly expressed than *SvAUX1* over all three time points. Among *SvAUX2-SvAUX5*, only *SvAUX4* differed significantly in *spp1* mutants, with higher expression in mutant plants than in A10.1 (Fig. 7B), possibly indicating a compensation effect. *SvAUX1, 2, 4*, and *5* belong to the turquoise module in A10.1, members of which are down-regulated at 10 DAS but up-regulated by 14 DAS. In *spp1* mutants, the expression pattern reverses for *SvAUX1* and *SvAUX2* (Fig. S11).

Only a few auxin-related genes differed significantly in expression between genotypes (Fig. 7C, Table S8). In A10.1 these fell into the turquoise, blue and brown modules (Fig. S11), which together include most of the transcripts. *SvVT2* and two auxin signaling F-box binding (AFB) genes (encoding potential auxin receptors), and a homolog of *BARREN STALK1/LAX PANICLE 1 (LAX1/SvBA1*, encoding a basic helix-loop-helix protein potentially involved in auxin signaling) (Komatsu et al., 2003; Gallavotti et al., 2004) were downregulated at 12 and/or 14 DAS (Fig. 7C). While five of the six AUX/IAA genes were up-regulated in the mutant, one (4G058700_AUX/IAA) was down-regulated (Fig. 7C, Table S8).

Genes whose homologs in maize are important for branch initiation and boundary formation were all downregulated in *spp1* (Fig. 7D, Table S7), including homologs of *TASSEL SHEATH1* (*TSH1*, encoding a GATA transcription factor (TF)) (Wang et al., 2009; Whipple et al., 2010), *BRANCH ANGLE DEFECTIVE1* (*BAD1*, a TCP TF) (Bai et al., 2012), *BARREN*

*STALK FASTIGIATE1* (*BAF1*, an AT-hook protein) (Gallavotti et al., 2011) and *LIGULELESS 1* (*LG1*, a nuclear localized protein) (Moreno et al., 1997; Lewis et al., 2014). Expression of homologs of the RAMOSA pathway genes *RA1* (encoding a Cys2-His2 zinc-finger TF) (Vollbrecht et al., 2005), *RA2* (a *LATERAL ORGAN BOUNDARY* (*LOB*) domain TF) (Moreno et al., 1997; Bortiri et al., 2006) and *RA3* (encoding a trehalose-phosphate phosphatase) (Satoh-Nagasawa et al., 2006), also was lower in *spp1* (Fig. 7D, Table S7). Because expression levels are standardized to reflect relative, rather than absolute, expression, the down-regulation is unlikely to reflect the lower number of branches in *spp1*.

In contrast, homologs of genes promoting meristem indeterminacy and inflorescence meristem identity were upregulated (Fig. 7D, Table S7), including *TAWAWA1* (*TAW1*) (Yoshida et al., 2013) and *TERMINAL FLOWER1* (*TFL1*) (Nakagawa et al., 2002; Danilevskaya et al., 2010; Hanano and Goto, 2011). A homolog of *ABERRANT PHYLLOTAXY 1* (*ABPH1*, a cytokinin-inducible type A response regulator), which controls phyllotactic patterning and meristem size (Lee et al., 2009), was also significantly upregulated in *spp1*(Fig. 7D, Table S7), as were homologs of *BROWN MIDRIB 1* and *3* (*SvBM1* and *SvBM3*)(Fig. 7D and Table S7). While BM1 and BM3 are involved in lignin synthesis in maize (Vignols et al., 1995; Halpin et al., 1998), they also affect kernel number, plant height, and days to flowering (Pedersen et al., 2005), traits associated with *spp1/aux1* mutations.

### SPP1/SvAUX1, but not the other four AUX1 homologs, is necessary for inflorescence branching

We used CRISPR-Cas9 technology with two guide RNAs to introduce mutations into all five putative auxin importers in accession ME034V, used for its high transformation efficiency (Zhu et al., 2017) (Fig. S12A). We obtained two independently edited *svaux1* single mutants, which exhibited a phenotype similar to that of the *spp1* mutant in the A10.1 background (Fig. S12B). We also retrieved two double mutants, *svaux1svaux5* (designated as *aux1,5* for short) and *svaux1svaux3* (*aux1,3*), one triple mutant, *svaux1svaux2svaux5* (*aux1,2,5*) and two quintuple mutants, *svaux1svaux2svaux3svaux4svaux5* (*aux1,2,3,4,5*) (Fig. S12B). One quintuple mutant, line cz66-11-16-11-1-4, had edits in all five homologs, with indels in *SvAUX2-SvAUX5* likely to knock out gene function because of frameshifts. However the edit in *SvAUX1* resulted in a single non-synonymous substitution (Fig. S12B), substituting an aliphatic residue (leucine) for an aromatic one (phenylalanine) in a presumed transmembrane domain (Fig. S6A); both residues are hydrophobic and will have limited effect on charge. We inferred that *SvAUX1* in this line could still be functional, leaving the line with only four mutated SvAUX1 homologs. Here we refer to this line as *svaux2svaux3svaux4svaux5* (*aux2,3,4,5*).

All *SvAUX* mutants were significantly shorter than wildtype after 4 weeks of growth, and all except *aux1,5* were still shorter than wildtype at 10 weeks, although leaf number was not significantly affected (Fig. 8A-8F, 8M; Table S9). Tiller number in wildtype plants did not differ between 4 and 10 weeks of growth, and the mutants did not differ amongst themselves at either stage (Fig. 8A-8F, 8N; Table S9). However, tiller number in the mutants was significantly higher than wildtype at 10 weeks despite having been the same at 4 weeks. Because *aux2,3,4,5* had more tillers, one of its the four mutant AUX loci likely contributes to the tillering phenotype in addition to SvAUX1 (Fig. 8A-8F, 8N).

**Figure 8.**
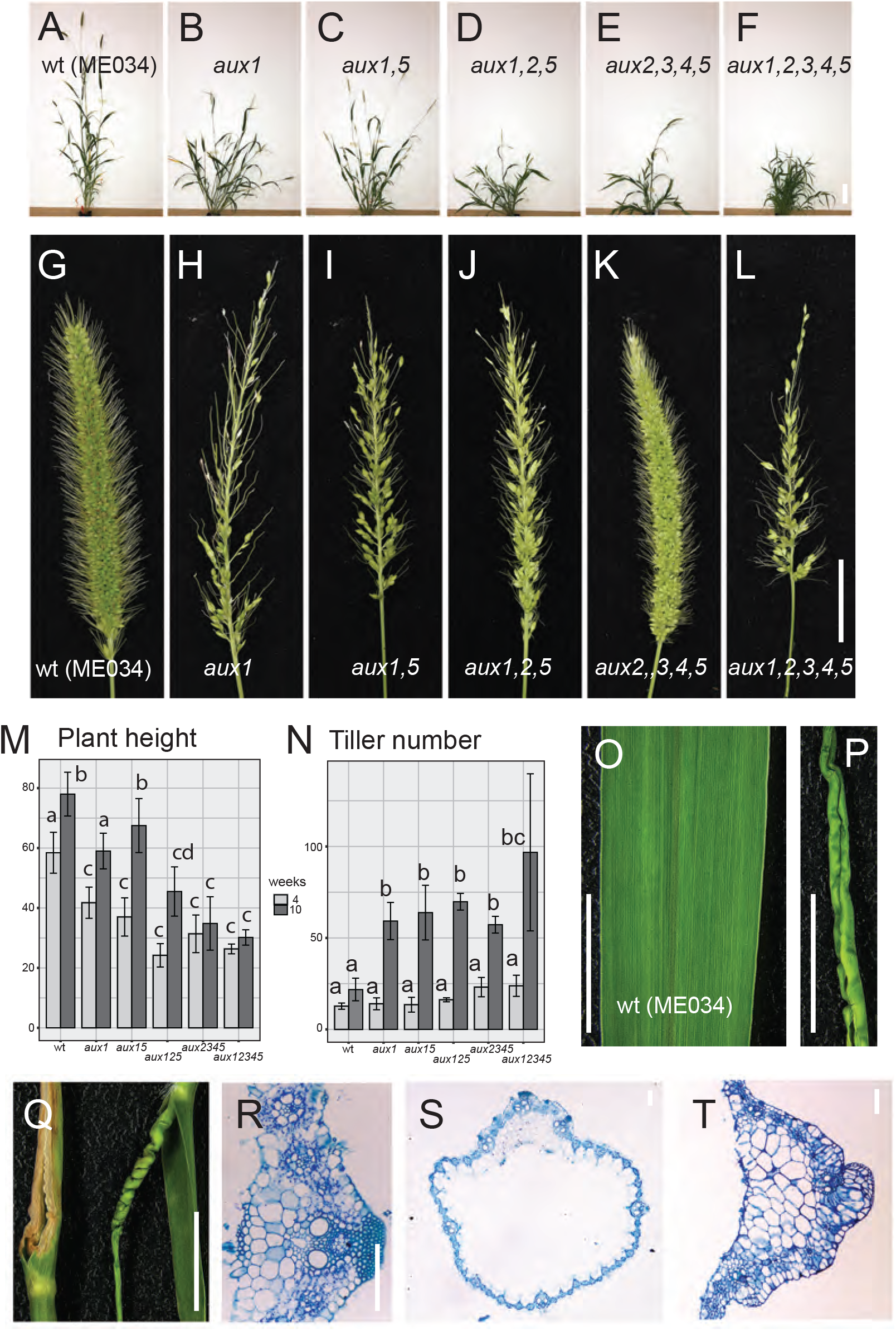
Auxin importer gene mutants in *S. viridis*. (A-F) wildtype and mutant plants photographed at 58 DAS, showing relative height and extent of tillering. (A) wildtype (ME034V); (B) *aux1*; (C) *aux1,5*; (D) *aux1,2,5*; (E) *aux2,3,4,5*; (F) *aux1,2,3,4,5. aux 1,3* not available for this set of photos. Scale = 10 cm. (G-L) wildtype and mutant inflorescences from the same plants and on the same day as in (A-F). Scale = 2 cm. (M) Plant height (mm) at the 4th (light gray) and 10th (dark gray) week after sowing. Error bars are standard deviations; values with the same letter are not significantly different by ANOVA. See also Table S10 for means, standard deviations, and p values. (N) Number of tillers on each plant at the 4th (light gray) and 10th (dark gray) week after sowing. Statistics as in (M). (O) wildtype ME034V (WT) leaf. Scale bar = 1 cm. (P, Q) Leaves in *aux1,2,5* or *aux1,2,3,4,5* mutants showing tube shape (P, right leaf in Q), early senescence in the tips (left leaf in Q), and twisted shape (right leaf in Q). Scale bar = 1 cm. (R-T) Leaf cross sections from WT (R), *aux1,2,5* (S) and *aux1,2,3,4,5* (T) mutants. Toluidine blue staining. Scale bar = 100 μm.

Inflorescences of higher order mutants involving *svaux1* were similar to those of *svaux1* single mutants (Fig. 8G-8L), consistent with our hypothesis that SvAUX1 is the major auxin influx carrier regulating inflorescence branching. Conversely, inflorescences of *aux2,3,4,5* were morphologically similar to those of wildtype (Fig. 8G, 8K), implying that the F377L substitution in SvAUX1 in that line indeed does not affect its function and that SvAUX1 alone is sufficient for inflorescence branch formation. Panicle length did not vary significantly among the plants, except that the panicle of *aux2,3,4,5* was slightly shorter, a difference that was just barely significant (Table S9).

The higher order mutants also exhibited phenotypes that were not observed in wildtype or single mutants (*svaux1* or *spp1*). For example, *aux1,2,5* and *aux1,2,3,4,5* often produced twisted or tube-shaped leaves, or leaves that senesced prematurely with yellowing tips and edges (Fig. 8O-8Q). Midrib cell layers and organization were affected in these mutants (Fig. 8R-8T). Lateral root number in the *aux1,5* and *aux1,2,5* was also reduced but primary root length was unaffected (Fig. S12C-S12E).

## DISCUSSION

The effect of AUX1 mutations on the inflorescence in *S. viridis* (*spp1/SvAUX1*) is strong, easily observed, and not obscured by mutations in its four paralogs, unlike mutations in AUX1 orthologs in other species such as Arabidopsis. The clear mutant phenotype has allowed us to uncover and validate numerous developmental roles for the auxin importer, including several that had not been observed in other systems. We were specifically interested in the role of SvAUX1 in inflorescence branching but also identified functions in stigma branching, formation of higher order inflorescence branches, and glumes (leaf-like floral bracts) which together affect plant fertility (yield). We extended our observations to maize, where, by manipulating other aspects of auxin synthesis, transport and signaling, we showed that ZmAUX1 also influences leaf number, which has not been observed previously. SvAUX1~GFP shows that the protein is membrane-localized and is expressed in inflorescence branch meristems, consistent with mutant analysis indicating that it is clearly necessary for inflorescence branching in *S. viridis*. Our data also add SPP1/AUX1 to the list of auxin transporters showing epidermal localization (Kubeš et al., 2012; Balzan et al., 2014; Swarup and Bhosale, 2019).

Of the five AUX1-like genes in *S. viridis*, SPP1/SvAUX1 has the major effect on inflorescence branching, although we cannot fully rule out the possibility that the other homologs could have a weak effect on their own. Consistent with this, AUX2 and AUX3 have low to no expression during inflorescence development. AUX4 and AUX5 are highly expressed during early inflorescence development, but mutations in these genes do not further enhance the sparse panicle phenotype of *spp1*; instead they lead to shorter plants. Assuming that the model of auxin flow in *S. viridis* is similar to that demonstrated in other species (e.g., O’Connor et al. (2014)), we speculate that AUX4 and AUX5 proteins could participate in internal basipetal auxin transport from auxin maxima at the branch initiation sites, whereas SPP1/AUX1 is likely mediating auxin movement to the branch initiation sites in the outer cell layers. Future imaging of the localization and dynamics of these auxin influx carriers is necessary to test this hypothesis.

Although SvAUX2-SvAUX5 make minimal or no contribution to inflorescence branching, they collectively are also important for plant height, tiller formation and leaf development. Reduced plant height and increased tiller number, as seen in higher order mutants, indicates a loss in apical dominance, a characteristic function of auxin. Twisted leaves are also seen in maize mutants whose auxin function is compromised, such as *growth regulating factor-interacting factor1* (Zhang et al., 2018) and *rough sheath 2* (Tsiantis et al., 1999).

### SPP1 regulates multiple aspects of inflorescence development downstream of meristem maintenance

The *spp1(svaux1)* mutant has fewer primary inflorescence branches, fewer higher order inflorescence branches, an altered ratio of bristles to spikelets, and defective stigmas, indicating that SPP1 controls branch initiation and elongation and meristem fate determination. The *zmaux1* mutant was also abnormal in these aspects, suggesting the role of SPP1 is likely conserved in the panicoid grasses. However, inflorescence meristem size is not affected in *spp1*, suggesting that SPP1 controls inflorescence development independent of meristem maintenance in grasses. This is consistent with findings from Arabidopsis, where the quadruple mutant of *aux1lax1lax2lax3* had a normal meristem, despite its defects in phyllotactic patterning (Bainbridge et al., 2008).

The only defective floral organ in *spp1* is the stigma, whereas other auxin-related grass mutants, such as *ba1* (Gallavotti et al., 2004) and *bif2* (McSteen and Hake, 2001), aborted multiple floral organs. Stigmas in most grasses are highly branched, and our data suggest that auxin transport is necessary for appropriate branch formation. Mutations in other genes such as those of a SHORT INTERNODES (SHI) family transcription factor (Yuo et al., 2012) also affect stigma morphology, suggesting that a specific network of genes regulating stigma formation remains to be discovered. The stigma defects could also contribute to reduced fertility in *spp1*, although auxin is also involved in fertilization and seed development (Robert et al., 2015; Figueiredo and Köhler, 2018), which remain to be investigated in *spp1*.

### SPP1 affects regulation of many branching-related genes, but not wholesale rewiring of the transcriptome

Several positive regulators of branch initiation, such as SvLG1 and SvTSH, are downregulated in *spp1* mutants. RA1-RA3 control meristem fate and determinacy and their expression was also altered in *spp1*. However, TSH4 (encoding a Squamosa Promoter Binding Protein TF) (Chuck et al., 2010; Whipple et al., 2010) is part of the same pathway as TSH1 and controls boundary formation during lateral organ initiation, but expression of TSH4 is unchanged in *spp1*. Some genes involved in auxin signaling and response are affected, including several (but not all) AUX/IAAs, ARFs and SvBA1, but they do not respond to the *spp1* mutation in a consistent manner, with some being upregulated and others downregulated.

Because *spp1* is a presumed transporter, effects on transcription must be indirect and are likely responding to levels of auxin. Even without active auxin import into the cell, it is still able to diffuse in but this is (presumably) a less tightly controlled process than transport. Thus the genes and processes that are downregulated are likely to be ones that require both rapid and precisely timed active transport.

We suggest that the *spp1/svaux1* mutants and the SPP1~GFP tagged line could provide useful tools with which to develop broader models of auxin flux into and out of cells. While most of the phenotypes we report are not unexpected for a protein that affects auxin, they show that auxin influx exerts a more central control over plant development than previously known. In particular, SPP1/SvAUX1 is clearly a central player in the genetic network that modulates all above-ground branching and could be used to test models of auxin regulation. Whether the effects we see in Setaria and maize indicate a fundamental difference between monocots and dicots in the role of auxin influx awaits testing in a broader set of species.

## MATERIALS AND METHODS

### Plant growth, phenotyping, and statistical comparisons

*Setaria viridis* accessions A10.1 and ME034V were grown in growth chamber and greenhouse conditions, respectively, following Acharya et al. (2017) and Zhu et al. (2018). The original *spp1* mutation was isolated from an A10.1 background; ME034V was chosen for CRISPR confirmation of the mutant phenotype because of its high transformation efficiency. Plant height, leaf number, panicle length, and branch number were measured as described in Huang et al. (2017) and Zhu et al. (2018). Fertility was measured as the ratio of spikelets with a fully developed upper floret to total spikelets; bristles were ignored for fertility measurements. Tillers were counted at 37 days after sowing (DAS) and plant height measured at 40 DAS. Histology and SEM followed Zhu et al. (2018). Inflorescence length, meristem width and height were measured using ImageJ (Schneider et al., 2012) from SEM photos.

For root phenotyping, sterilized seeds were grown either in Murashige and Skoog (MS) medium or germination pouches as described in Huang et al. (2017) and Acharya et al. (2017), respectively.

Auxin rescue experiments followed Marchant (1999) and Yu et al. (2015). 2, 4-D (from Plant Media, in 1mM stock with pure ethanol) and NAA (from Sigma-Aldrich, in 10mM stock with pure ethanol) were added to the medium to a final concentration of 0.1 mM. MS medium containing 0.1% ethanol was used as a mock control. Seeds were grown on MS medium for three days and then transferred to media containing appropriate concentrations of auxin or mock for three more days. Root hairs were imaged at 4x magnification on a Leica DM750 microscope. Root hair number was counted in the focal plane on the side of the root facing the observer and normalized to root length. Experiments were repeated three times.

In maize, *zmaux1* mutant plants were crossed to *vt2, bif2* and *Bif4* mutants, and F2 segregating populations were grown in the field in Columbia, Missouri in 2017. Plants were genotyped to identify single and double mutants using primers listed in Table S10 and were phenotyped at the eighth week. For the dominant mutant *Bif4*, both heterozygotes and homozygotes were included for mutant phenotyping analysis. For each mutant and mutant combination we assessed traits of the tassel (length from flag leaf to tassel tip, number of branches, spikelets on main spike, spikelet number per cm) and ear (kernel number, ear row number), and three vegetative traits (height of flag leaf, number of leaves above the lowest elongated internode, and number of tillers).

All pairwise comparisons used Welch’s t-test as implemented in R (R Core Team, 2020). Single, double and higher order mutants were compared to each other and to wildtype by one-way or two-way Type I or Type II ANOVA as appropriate, followed by Tukey’s HSD test using standard programs in R (R Core Team, 2020). Comparisons with p>0.05 were considered non-significant.

### Generation of mutants for auxin importer gene homologs

Cloning of CRISPR-Cas9 constructs and *S. viridi*s transformation followed Zhu et al. (2018). Two guide RNAs targeting GGGAGATCATGCACGCGATG and AGTTGATGGGCCCGAAGAAG, respectively, were designed to target all five *SvAUX1* paralogs and used in the CRISPR-Cas9 constructs, which were then transformed into the accession ME034V. More than ten transgenic plants were obtained and gene edits in the auxin importer genes were examined using primers listed in Table S10. Stable homozygous lines in T3 or T4 were used for phenotypic analysis.

### RNAseq sampling, sequencing and analysis

Inflorescences from both A10.1 control plants and *spp1* mutants (in the A10.1 background) were dissected at 10, 12, and 14 days after sowing (DAS) for RNA extraction and library preparation with three to four biological replicates per genotype and stage, following Zhu et al. (2018). 100 bp paired-end sequences were produced on the Illumina HiSeq 2500 platform at the University of Illinois at Urbana-Champaign W.M. Keck Center.

Adaptors and low-quality reads were trimmed using Trimmomatic (Bolger et al., 2014) and reads were quality-checked using fastqc after trimming. The *S. viridis* reference genome (v1) was indexed using bowtie 2 (Langmead and Salzberg, 2012) from Sviridis_311_v1.0.fa.gz file at PhytozomeV11 (phytozome.jgi.doe.gov). Reads were mapped to the reference genome using tophat2 and differentially expressed genes were identified using cuffdiff (Trapnell et al., 2012). Expression levels quantified in Fragments Per Kilobase of transcript per Million (FPKM) were extracted for 35,214 *S. viridis* primary transcripts (Tables S5, S6). Gene annotation and grass homolog identification followed Zhu et al. ((2018)).

Genes with an average FPKM ≥5 per sample group (3-4 biological replicates) were extracted, and the log_2_(FPKM+1) of genes within the top 75% of the highest median absolute deviation (MAD) across three developmental stages was selected for co-expression analysis (nGenes = 10,434 from both genotypes). A co-expression network was constructed for each genotype using the R package WGCNA (v1.70) (Langfelder and Horvath, 2008) with an established pipeline (Yu et al., 2020), with blockwiseModules function and the following parameters: soft-thresholding power of 18, minModuleSize of 100, detectCutHeight of 0.995, mergeCutHeight of 0.25, deepSplit of 2. Degree distributions in each individual network followed the power law and satisfied the scale-free topology. Conservation of modules was tested with the modulePreservation function in the WGCNA package (Langfelder et al., 2011) following Yu et al. (2020). An improved *S. viridis* gene ortholog (GO) annotation was generated by the GOMAP annotation pipeline (Wimalanathan and Lawrence-Dill, 2021). 33391 of 35214 genes (representing 94.8% of primary transcripts in the *S. viridis* genome v1.1) were successfully annotated, with the median number of annotation terms per gene of 8. GO enrichment analysis and visualization used the R package clusterProfiler (v4.0) (Wu et al., 2021). The chord diagram of changes in module membership was plotted with R package circlize (v 0.4.13).

### Creation of SvSPP1~iGFP fusion protein, subcellular localization, and transgenics

Binary vectors were built using standard Golden Gate assembly (Werner et al., 2012). *SPP1* was internally tagged (hereafter, *SPP1~iGFP*) and placed either under the native *SvSPP1* (*proSvSPP1::SPP1-iGFP*) or a constitutive *Panicum virgatum UBI1* promoter (*proPvUBI1::SPP1-iGFP*). We were unable to transform *S. viridis* with the C-terminal fusion of GFP (SvSPP1-GFP), a problem also encountered in Arabidopsis by Swarup et al. (2004) for C- and N-terminal reporter fusions of auxin influx carriers including AtAUX1. Hence, we chose an internal facing (cytoplasmic) N-terminal hydrophilic loop of SPP1 because a similar AtAUX1 construct retained its topology and physiological role (Swarup et al., 2004). The GFP sequence in SvSPP1~iGFP was inserted between Lys_121_ and Asn_122_ (Fig. S6A), predicted to be in a hydrophilic loop (Swarup et al., 2004). 3kb of *SvSPP1* upstream sequence was PCR-amplified using genomic DNA and used as *proSvSPP1. SPP1a* (1-363), *SPP1b* (363-1470) and *GFP* fragments were PCR-amplified using either cDNA or the plasmid *pL0M-C2-eGFP-15095* as templates; primers are listed in Table S10. Each PCR fragment was cloned individually into the level 0 vectors *pICH41233* (*proSvSPP1*), *pICH41258* (*SPP1a*), *pAGM1299* (*GFP*), and *pAGM1301* (*SPP1b*). The resultant level 0 constructs plus level 0 NosT (*Nopaline synthase* terminator) vector were subsequently cloned in the level 1 vector *pICH47742* to produce *pICH47742-proSvSPP1::SvSPP1-iGFP::NosT*. The Level 1 construct *pICH47802*-*proZmUBI1::HPT*, an expression cassette with a functional *HPT* (*hygromycin phosphotransferase*) gene under a constitutive *Zea mays UBIQUITIN 1* promoter (*proZmUBI1::HPT*), and the *pICH47742-proSvSPP1::SvSPP1-iGFP::NosT* were then assembled in the binary level 2 vector *pICSL4723*.

The binary vector was transformed in *Agrobacterium tumefaciens* strain AGL1 for transient (tobacco) or transgenic (*S. viridis*) expression analysis. To check transient expression, six-week-old *Nicotiana benthamiana* leaves were agro-infiltrated as previously described (Cho et al., 2015). After 4 days, GFP fluorescence was visualized using a Leica SP8 (USA) confocal laser scanning microscope. Excitation and emission wavelengths for GFP and chlorophyll were 488/ 510-540 nm and 561/ 673-726 nm, respectively.

The binary vector was stably transformed into the *spp1-1* mutant line (Huang et al., 2017) at the Donald Danforth Plant Science Center Plant Transformation Facility (St Louis, MO). Five putatively transgenic plants were obtained and presence of GFP was confirmed in three of them using PCR genotyping with GFP-specific primers in the T_0_ generation. One line homozygous for the transgene (GFP) was chosen and its stable expression was used in subsequent generations for confocal imaging (T_3_) and phenotypic analysis (T_4_). Primers for genotyping and expression assays are listed in Table S10.

Transgene expression was validated by RT-qPCR. 4DAS leaves (3^rd^ leaf base) and 11DAS primary inflorescences were hand-dissected as described in Li et al. (2010) and Huang et al. (2017), respectively. Four or five plants were pooled for each biological replicate. Total RNA was extracted using an RNeasy Plant Mini Kit (Qiagen) and quantified using a NanoDrop 1000 spectrophotometer (Thermo-Fisher). Each RNA sample was reverse-transcribed to cDNA after DNase I treatment using a PrimeScript RT reagent kit (Takara). PCR was performed as described in Kumar et al. (2017). *Sevir*.*9G574400* and *Sevir*.*2G354200* were used as reference genes as described in Huang et al., (2017). The normalized relative quantity of GFP transgene to the two reference genes was estimated using the Comparative CT Method (ΔΔ^CT^ method) (Schmittgen and Livak, 2008).

### Image capture, analysis, and processing

Confocal images were captured on a Leica TCS SP8 confocal laser scanning microscope with an HC PL APO CS2 63x, 40x and 20x /1.20 WATER objective lens (Leica Microsystems, Mannheim, Germany) and Leica Application Suite X (LAS X) software. The light source was the White Light Laser (WLL for GFP, chlorophyll, and FM4-64), while emission fluorescence was captured by the hybrid (HyD™) detector. Excitation and emission wavelengths for GFP, FM4-64 and chlorophyll were 430/480 nm, 490/550 nm, and 561/673-726 nm, respectively. For bright field images, a conventional photomultiplier tube (PMT) for transmittance was used (PMT trans in LAS X software). For image capture, line averages and frame accumulations were 6-16 (for roots) and 3-12 times (for inflorescence and leaves) to reduce noise. Inflorescence meristems and leaf cross sections were imaged as Z-stacks; images were reconstructed using Imaris x64, 7.2.3 (www.bitplane.com) with background subtraction settings enabled. SPP1 cellular localization in transgenic tissues was observed through multiple confocal sections. Four or five inflorescences from 11DAS plants were dissected under the stereomicroscope and analyzed. The fourth leaf from the base from 6DAS plants was embedded in 6% agarose, sectioned using a Vibratome (1500 Sectioning System), and stained using FM4-64 as described by Grandjean et al. (2004) before imaging.

All images in this paper were resized as necessary, adjusted for brightness, and assembled into figures in Adobe Photoshop. Images were then imported into Adobe Illustrator for labeling. Graphs were produced with ggplot2 in R and also imported into Illustrator to adjust labels and line width.

## ACKNOWLEDGEMENTS

We thank Daniel Voytas (University of Minnesota) for sending us pMOD_A1110 (pJG471), pMOD_B2518 (pJG310), pMOD_C2616 (pJG338) and pTRANS_250d (pRLG103) constructs for CRISPR-Cas9 cloning. Taylor AuBuchon-Elder, Michael Cambron and Callista Martin provided technical assistance early in the project. We thank Howard Berg and Kirk Czymmerk of the Advanced Bioimaging Laboratory at the Danforth Center for their technical help with confocal imaging and data analysis.

## DATA AVAILABILITY

Raw sequence reads for RNA-seq in A10.1 through development are deposited at the NCBI Gene Expression Omnibus (GEO) under the accession number GSE118673 (Zhu et al. (2018). Reads for the *spp1* mutant are in GEO under number XXX (to be inserted after manuscript acceptance). Raw phenotype data are in datadryad accession number XXX (to be inserted after manuscript acceptance).

**Figure S1. Additional phenotypes of *spp1* mutant plants**. (A-G) Shoot phenotypes comparing wildtype (A10.1, white bars) with *spp1* mutants (black bars). (A) Days to heading. (B) Days to anthesis. (C) Size of upper (fertile) floret (mm). (D) Percent seed germination. (E) Peduncle diameter. (F, G) Cross sections of peduncles stained with toluidine blue. (F) wildtype (A10.1); (G) *spp1*. Arrows, vascular bundles; scale bar = 200 μm. (H-N) Root phenotypes comparing wildtype (A10.1, white bars) with *spp1* mutants (black bars). (H, I) Density of root hairs on the main root (H) and lateral roots (I). (J, K). Number of root hair initials on the main root (J) and lateral roots (K). (L, M) Main and lateral roots of wildtype (L) and *spp1* (M) showing differences in root hair density. Scale =2 mm. (N) Washed root systems of wildtype (left) and *spp1* (right) showing similar sizes. Scale = 1 cm. Error bars are standard deviations. Significance values determined by Welch’s t-test. = 0.01-0.05, *, <0.01, **, <0.001, ***, <0.0001.

**Figure S2. Auxin rescue experiments**. (A-F) Root growth and gravitropism of A10.1 (A, C and E) and *spp1* (B, D and F) at mock (A and B), 0.1 μm 2,4-D (C and D) and 0.1 μm NAA (E and F) treatments. Scale bar = 3 cm. (G-L) Root hairs of A10.1 (G, I and K) and *spp1* (H, J and L) at mock (G and H), 0.1 μm 2,4-D (I and J) and 0.1 μm NAA (K and L) treatments. Scale bar = 1 mm. (M) Root hair density on the primary roots in A10.1 and *spp1* with different auxin treatments. Significance assessed by ANOVA and Tukey’s HSD. See Table S1 for means, standard deviations, and p values.

**Figure S3. Phenotype of *zmaux1vt2* double mutants**. (A) Representative whole plant pictures. (B) Ear row number. (C) Spikelets per row. (D) Total number of kernels. (E) Number of tassel branches. (F) Number of spikelets on the main spike of the tassel. (G) Number of spikelets per cm (spikelet density). (H) Tassel length (cm). (I) Flag leaf height (cm) from ground. (J) Total number of leaves. Branch number, tassel spikelet number per cm and kernel number measured at 56 DAS. Left to right, WT (white bars), *zmaux1* (gray bars), *vt2* (gray bars), *zmaux1vt2* (black bars). Error bars are ± one standard deviation. Values with the same letter are not significantly different at p =< 0.05. Significance assessed by ANOVA and Tukey’s HSD. See Table S3 for sample sizes, means, standard deviations, and p values.

**Figure S4. Phenotype of *zmaux1bif2* double mutants**. (A) Representative whole plant pictures. (B) Ear row number. (C) Spikelets per row. (D) Total number of kernels. (E) Number of tassel branches. (F) Number of spikelets on the main spike of the tassel. (G) Number of spikelets per cm (spikelet density). (H) Tassel length (cm). (I) Flag leaf height (cm) from ground. (J) Total number of leaves. Branch number, tassel spikelet number per cm and kernel number measured at 56 DAS. Left to right, WT (white bars), *zmaux1* (gray bars), *bif2* (gray bars), *zmaux1bif2* (black bars). Statistics as in Figure S3.

**Figure S5. Phenotype of *zmaux1Bif4* double mutants**. (A) Representative whole plant pictures. (B) Ear row number. (C) Spikelets per row. (D) Total number of kernels. (E) Number of tassel branches. (F) Number of spikelets on the main spike of the tassel. (G) Number of spikelets per cm (spikelet density). (H) Tassel length (cm). (I) Flag leaf height (cm) from ground. (J) Total number of leaves. Branch number, tassel spikelet number per cm and kernel number measured at 56 DAS. Left to right, WT (white bars), *zmaux1* (gray bars), *Bif4* (gray bars), *zmaux1Bif4* (black bars). Statistics as in Figure S3.

**Figure S6. Cellular localization of SPP1~iGFP**. (A) Schematic diagram of SPP1 protein topology showing hydrophilic regions predicted to be in extra- and intracellular spaces. Green arrow indicates the position of GFP inserted in the N-terminal cytoplasmic loop (internal) to test SPP1~iGFP localization. Closed blue circle indicates the position of Phe_377_ to Leu_377_ substitution in the *svaux1* gene of *aux1,2,3,4,5*. (B-D) Confocal images of tobacco leaf cells transiently expressing SPP1~iGFP, showing localization to a thin line around the cell. Panels from left to right: SPP1~iGFP (B), chlorophyll autofluorescence (C), and overlay (D). Scale = 20 μm. (E-J) Stable expression of SPP1~iGFP in roots of *S. viridis* at 9 DAS. Imaging of root tissues focused on either inner (E, F) or outer tissues (G-J) showing fluorescent signals on the plasma membrane (PM), predominantly in the epidermis. (I) Enlarged image of the boxed region of (H), confirming GFP signals around the nuclear membrane (yellow arrowheads). (J) Non-transgenic control.

**Figure S7. Validation of SPP1~iGFP in transgenic *S. viridis***. (A) Gel image of PCR results confirming the presence of GFP band (~188bp, bottom bands) in transgenic *S. viridis* plants. PCR bands at ~540bp correspond to an *S. viridis* gene (*Sevir*.*2G209800*) serving as a positive control. (B) RT-qPCR assay determining the expression of SPP1~iGFP in transgenics. Relative expression was quantified for *GFP* in leaf at 4 DAS (3^rd^ leaf base; N = 4, pooled) and dissected inflorescence primordia at 11 DAS (N = 5, pooled), respectively. Data are the mean of three technical replicates. Expression data for *GFP* were normalized to expression of reference genes *Sevir*.*2G354200* and *Sevir*.*9G574400*. (C-K) Expression of SPP1~iGFP partially rescued the *spp1* defects in inflorescence and roots. (C) Representative plants from A10.1, non-transgenic (*spp1_NT*) and transgenic (*spp1_T)* lines at 26 DAS. (D) Plant heights at 23, 34, and 40 DAS for the three genotypes. (E) Representative panicles from A10.1, non-transgenic (*spp1_NT*) and transgenic (*spp1_T)* plants at 30 DAS. (F-I) Inflorescence traits for all three genotypes at 35 DAS. (F) Panicle length. (G) Primary branch number. (H) Spikelet number per branch. (I) Bristle number per branch. (J) Root growth assay showing agravitropic response of *spp1_T* seedlings at 5 DAS. (K) Numbers of agravitropic seedlings in wt and transgenics. Bars represent mean values, error bars show standard deviations; data summarized in Table S4. Statistics as in Figure S3.

**Figure S8. Gene co-expression modules**. Weighted gene correlation network analysis (WGCNA) detected seven co-expression modules in wild *S. viridis* A10.1. (A) Cluster dendrogram shows co-expression module assignment. (B) Expression patterns of module genes and module eigengene are shown by heatmap (top) and bar graph (bottom), respectively.

**Figure S9. Comparisons between wild *S. viridis* A10.1 and *spp1* mutant networks**. (A) Preservation analysis of WGCNA modules in the reference genotype (wildtype *S. viridis* A10.1) versus the test genotype (*spp1* mutant) and conversely. (B). Zsummary>10, high preservation, 2< Zsummary<10, weak to moderate preservation, Zsummary<2, no preservation. (C) Similarity analysis using numbers of overlapping genes in WGCNA modules between genotypes, showing the number of overlapping genes and p-values from Fisher’s exact test (in parentheses). White to red color gradient indicates –log10(p-value).

**Figure S10. GO enrichment**. GO enrichment analysis of major WGCNA modules in *S. viridis* A10.1, and *spp1* mutant. Dot color represents statistical significance of the enrichment (adjusted p-value, a color gradient from blue (<0.01) to red (<0.05)). The sizes of the dots represent gene ratio (number of significant genes/number of annotated genes in each GO term). GO terms were not significantly enriched in red and grey modules in A10.1, and grey module in *spp1* (not displayed).

**Figure S11. Chord diagram illustrating how WGCNA module membership differs between genotypes**. Color keys on the left side of the diagram represent the seven modules identified by WGCNA in wild *S. viridis* A10.1, and on the right side represent the ten modules in *spp1*. Paths of reassignment of genes are illustrated as flows in the diagram. The inner color keys ring of wild *S. viridis* A10.1 (left half) represents the reassigned modules in the *spp1* mutant. The three dashed lines show changes of module membership of five auxin-related genes. **Figure S12. Mutants of auxin importer genes**. (A) Target sequences for gRNA1 (cyan arrow) and gRNA2 (magenta arrow), respectively. Boldface letters represent the PAM sites. On the gene models for the five auxin importer genes in *S*.*viridis*, SvAUX1-SvAUX5, cyan and magenta arrows show locations of target sites. Numbers at ends of arrows indicate number of mismatches between gRNA and target sites. (B) Table of edits at each gRNA target site in each of the five auxin influx carrier genes in each line. x, no editing; +, addition; -, deletion; bp, base pair; ->, substitution. (C) Roots of ME034V, *aux1,5* and *aux1,2,5* in 7 DAS plants. (D) Lateral root number and (E) primary root length in these plants.

**Video S1**. Confocal 3D reconstruction of a single inflorescence meristem corresponding to that shown in Figure 6G for Setaria SPP1~GFP expression domains.

**Video S2**. Confocal 3D reconstruction of early development of a single inflorescence corresponding to that shown in Figure 6H for Setaria SPP1~GFP expression domains. Image shows primary and some secondary branch meristems.

## List of supplemental tables with brief titles

**Table S1. Phenotypic comparisons between A10.1 and *spp1* mutants**.

**Table S2. Phenotypic comparisons between A10.1 and *spp1* mutants over development**.

**Table S3. Phenotypic comparisons between W22 (maize wildtype), *zmaux1* and single and double mutants of selected genes in the auxin pathway**.

**Table S4. Phenotypic comparisons between A10.1, spp1_T and spp1_NT, testing for complementation of SPP1~GFP**.

**Table S5. RNA-seq library sequencing and mapping statistics**.

**Table S6. Expression of all *S. viridis* genes from each replicate (R1-R4) of the developmental stages (10, 12, 14 DAS) in A10.1 and *spp1***.

**Table S7. Expression of differentially expressed genes between A10.1 and *spp1* at each developmental stage (10, 12, 14 DAS)**.

**Table S8. Expression of auxin pathway related genes in A10.1 and *spp1* at each developmental stage (10, 12, 14 DAS)**.

**Table S9. Phenotypic comparisons among AUX CRISPR mutants.**

**Table S10: Primers used in this study**.

## Notes

1 C.Z., M.S.B. and D.T. were supported by National Science Foundation (NSF) grant IOS-1413824 to E.A.K.

